# Porthole and Stormcloud: Tools for visualisation of spatiotemporal M/EEG statistics

**DOI:** 10.1101/534784

**Authors:** Jeremy A Taylor, Marta I Garrido

## Abstract

Electro- and magneto-encephalography are functional neuroimaging modalities characterised by their ability to quantify dynamic spatiotemporal activity within the brain. However, the visualisation techniques currently used to illustrate these effects are currently limited to single- or multi-channel time series plots, topographic scalp maps and orthographic cross-sections of the spatiotemporal data structure. Whilst these methods each have their own strength and weaknesses, they are only able to show a subset of the data and are suboptimal at articulating one or both of the space-time components.

Here, we propose *Porthole* and *Stormcloud*, a set of data visualisation tools which can automatically generate context appropriate graphics for both print and screen with the following graphical capabilities:

1. Animated two-dimensional scalp maps with dynamic timeline annotation and optional user interaction;
2. Three-dimensional construction of discrete clusters within sparse spatiotemporal volumes, rendered with ‘cloud-like’ appearance and augmented by cross-sectional scalp maps indicating local maxima.

These publicly available tools were designed specifically for visualisation of M/EEG spatiotemporal statistical maps, however, we also demonstrate alternate use cases of posterior probability maps and weight maps produced by machine learning classifiers. In principle, the methods employed here are transferrable to visualisation of any spatiotemporal image.

## 1. Introduction

The ability to effectively communicate experimental findings is central to any scientific endeavour. As the complexity of our research increases, there is an inherent tradeoff between the time investment necessary for the author to explain their ideas and for the audience to understand them (Olah and Carter, 2017). Data visualisation is a critical, yet often overlooked mode of communication with the potential to bridge this gap by accentuating or summarising the author’s key messages in a clear and efficient manner. However, whilst there are a number of generic visualisation tropes that the reader has grown accustomed to, we suggest that these are largely suboptimal at articulating the necessary context surrounding the data in some domain-specific subfields of neuroscience.

Electroencephalography (EEG) and magnetoencephalography (MEG) are functional neuroimaging modalities which measure fluctuations in electrical and magnetic components of the electromagnetic field, respectively, as generated by the brain in action. Both methods are characterised by an excellent temporal resolution and a high density of sensors or *channels*, often providing whole-scalp coverage. Whilst M/EEG data can be analysed on a discrete single-channel basis (Figure 1A), these channels can also be interpolated to create a two-dimensional map (Figure 1C) of brain activity over the full surface of the scalp (Koles and Paranjape, 1988). As such, the time series data from these recordings can be processed as a three-dimensional volume, comprising the two spatial dimensions on the surface of the scalp over time. We refer to these volumes as *spatiotemporal* or *scalp-time* images. In terms of analysis, transforming the data in this manner is particularly useful for computational modelling, enabling us to obtain spatiotemporal statistical parametric maps (SPM; Friston et al., 2011; Litvak et al., 2011) and inquire about regionally specific statistical effects across the dataset as a whole, without bias or making any a priori assumptions or bias, i.e. pre-selecting a subset of the data, such as specific channels and/or time components of the event-related potential/field (ERP/ERF).

Spatiotemporal images are by no means a new construct, nor specific to neuroscience. The generalist three-dimensional space model, representing a two-dimensional space over time is more commonly referred to as a *space-time cube*, originally devised by Torsten Hägerstrand (1970) for analysing social interactions on a geographical map. This model has since been applied to many other domains, each with their own specific visualisation challenges and solutions, enabling the most important information within the volume to be easily interpretable by the intended audience. These challenges primarily stem from the fact that by presenting three-dimensional data on a two-dimensional medium, whether this be in print or on screen, we are only able to accurately represent a subset of the data. These two-dimensional visualisations can all be expressed in terms of the *operations* which are applied to the space-time cube to extract these data. For a comprehensive review of these operations, refer to Bach et al. (2016). Here, we critique the core visualisation methods currently being used for presenting M/EEG analysis; single- or multi-channel plots, serial topographic scalp maps and orthographic projections, and later propose alternatives that we believe capture more information from the data.

A single-channel time series (as shown in Figure 1A) can be described as a vector extracted from a single point on the spatial *x*-*y* plane and parallel to the *t-* axis, an operation referred to as *time drilling* (Bach et al., 2016). The responses measured simultaneously from all sensors can also be overlaid onto a multi-channel or ‘butterfly’ plot (Figure 1B, or *repeated drilling* in multiple spatial locations). Whilst this representation of the data is useful for understanding how the signal recorded from one channel evolves over time, it provides no information on the spatial profile.

Another conventional way of displaying M/EEG data is through topographic scalp maps (Figure 1C), which typically display ERP/ERF activity spatially interpolated over channels at a given time point, a range of time points, or the average over all time points. In other words, this data representation can be obtained by slicing a spatiotemporal volume of data across the *x*-*y* plane for a range of time points, *t*, a process otherwise referred to as *time cutting* (Bach et al., 2016).

**Figure 1.**
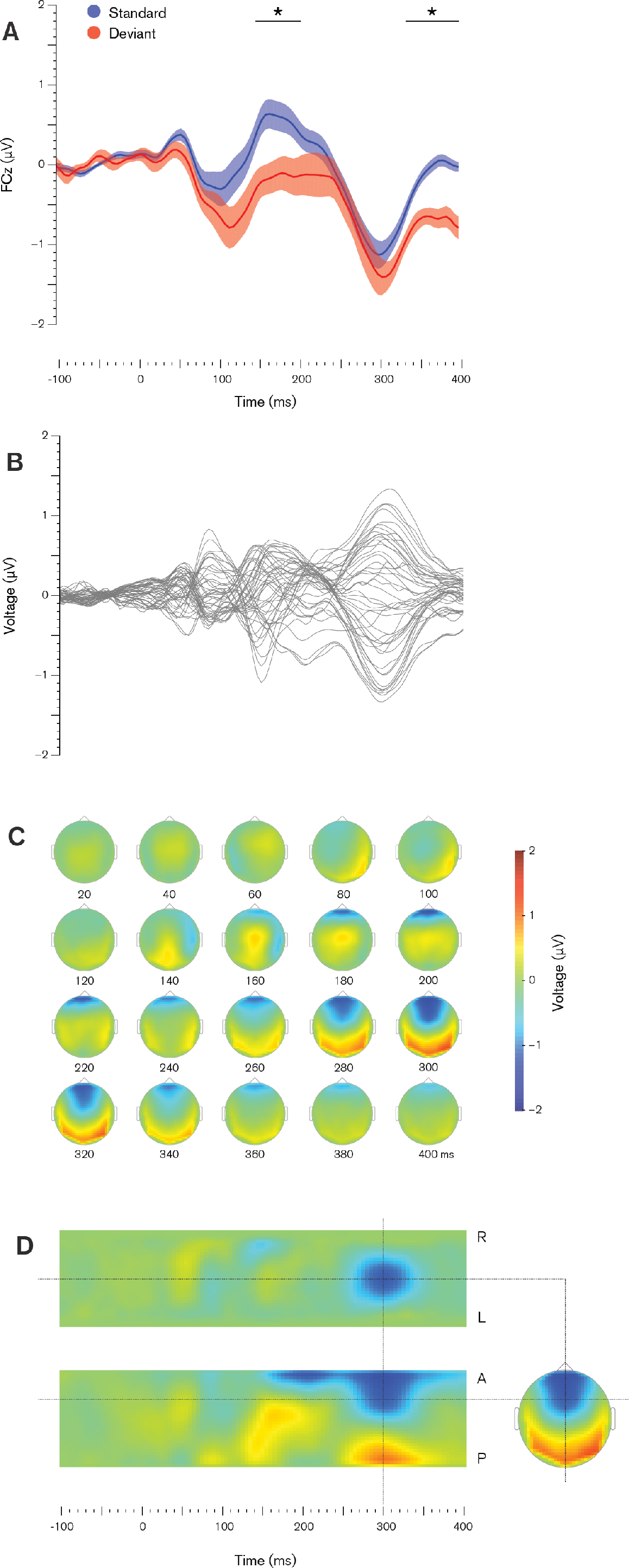
Examples of existing M/EEG visualisation methods. **(a)** *Time drilling* — Single channel plot of grand-average brain responses evoked by attended standard (blue) and deviant (red) stimuli, recorded from central (Cz) channel. Standard error shown as shaded areas, star indicates time points of significant different responses (*p* < 0.05, paired *t*-test, FDR corrected). **(b)** *Repeated drilling* — Butterfly plot of grand-average response to attended deviant stimuli. Each line indicates the ERP measured at one of 64 channels. **(c)** *Time cutting* — Topographic scalp maps of grand-average response to attended deviant stimulus, sampled at 20 ms intervals from 0 to 400 ms. **(d)** *Space cutting* — Orthographic projection of grand-average response to attended deviant stimulus, sectioned through the Cz channel at time 250 ms.

Topographic scalp maps are often presented at specific time points of interest, or as a sequence at set intervals over the length of the *t*-axis, akin to small multiples, described by Tufte (2001) as a set of small figures on shared axes for direct comparison. However, as the amount of paper required to display the full sequence of scalp maps at the native sampling frequency is prohibitive, typically only a subset of these time points are shown. As such, presenting the data in this manner is able to provide high spatial resolution, but low temporal resolution.

An orthographic projection (Figure 1D) comprises a set of three planar cross-sections which intersect orthogonally through a single point within the spatiotemporal volume, denoted here as (*x*,*y*,*t*). In addition to a topographic scalp map (i.e. cutting across the *x*-*y* plane at time *t*), temporal slices are made perpendicular to both spatial axes in the *x*-*t* and *y*-*t* planes, an operation referred to as *space cutting* (Bach et al., 2016). These projections offer high temporal resolution and moderate spatial resolution with respect to the point of interest but mask all other information as a result. Such operations may be useful when the volume can be navigated by moving the point of interest via user interaction, but are less than ideal for a static two-dimensional medium.

Although M/EEG modalities are able to capture spatial and temporal changes in brain activity, all of these visualisation techniques currently being employed are limited in their ability to articulate one, but not all space-time components of the data. Over-reliance on these methods may be due to a dearth of freely available user-friendly tools which can be used by both programmers and non-programmers alike, designed specifically for these modalities and spatiotemporal data structures.

In this paper, we introduce *Porthole* and *Stormcloud*, a set of visualisation tools designed to address these shortcomings, demonstrating improvements in the illustration of sparse spatiotemporal data through interactive two-dimensional animations and three-dimensional rendering. In the Methods section, we provide a summary of the data structure, break down each individual component of the visual environment, and outline the operations we perform in generating the visualisations.

In the Results section, we provide examples of figures produced when applying these tools to outputs from three different analyses on empirical data; namely statistical parametric maps, posterior probability maps, and machine learning weight maps. Finally, we discuss the key advantages, limitations, and possible future directions for this work.

## 2. Methods

### 2.1 Programming language

We chose to write these tools for MATLAB (The Mathworks Inc., Natick, Massachusetts), given that it is currently the primary software used by the neuroimaging community. In this way, the user does not need to leave their existing workflow or install additional software. This also avoids potential compatibility issues, as any operating system and hardware configuration currently running MATLAB should be able to use these visualisation tools.

### 2.2 Data model

The visualisation methods employed here were developed specifically for use with scalp-time images generated using Statistical Parametric Mapping (SPM; Friston et al., 2011), a univariate modelling technique commonly used for analysing neuroimaging data, as implemented within a freely available software package (Litvak et al., 2011). In Section 2.2.1, we provide a brief overview of the SPM operations and terminology, the types of images produced and the effects of interest within these images. Using this framework as a basis allowed us to make some reasonable assumptions about the data, as outlined in Section 2.2.2, and directly informed our design approach, detailed in Sections 2.3 and 2.4.

#### 2.2.1 Statistical parametric mapping

SPM assumes that the distribution of values contained within the image voxels are members of a known probability density function, namely Student’s *t* or *F* distributions with mean zero. Any values considered unlikely to be drawn from this distribution are interpreted as effects resultant from the experimental conditions.

To make such inferences, a mass-univariate approach is employed by independently estimating general linear models (GLM) at each individual voxel by regressing all of the experimental variables across all participants. The resulting coefficients, or *β* parameters from these models represent the proportion of signal that can be explained by the experimental conditions (Friston et al., 2011). To test hypotheses about these conditions, *t-* or *F*-statistics and their associated *p*-values are obtained for each voxel as a linear combination of the *β* parameters and residual variance, also known as a *contrast* (Poline et al., 2007). *t-* or *F*-contrasts differ in the types of questions they can answer: the SPM *t*-test uses a single-tail and is only able to test for positive effects (whether the response to one condition is greater than another), whereas the *F*-test is unsigned and can test for both positive and negative effects (whether the responses to conditions differ).

We therefore arrive at a corresponding three-dimensional map of *t*- or *F*-values and wish to consider the volume as a whole. However, as the spatiotemporal volume can comprise in the order of 100,000 voxels, it becomes increasingly likely to make false discoveries when performing a large number of statistical tests simultaneously. To determine where the effects of interest are localised, the statistic map is thresholded at a given level of confidence, and any groups of voxels, or *clusters* of activity above this threshold are examined. This threshold level can be computed using random field theory and family-wise error (FWE) correction for multiple comparisons over all voxels (Brett et al., 2004; Worsley, 1995, 1996), or remain uncorrected at the discretion of the user. We can then draw conclusions about this thresholded data at the *cluster*-level (the chance of finding another cluster containing this number of voxels) or at the *peak*-level (the chance of finding another single voxel of this value). Below, propose techniques to visualise these three-dimensional *t*-/*F*-maps in a way which preserves both the higher temporal resolution of the data and the spatial topology of the activation clusters.

#### 2.2.2 Data characteristics

Using this SPM framework as a basis, we are able to make a number of key reasonable assumptions about the data and use these to directly inform our design approach:

1. **Sparsity.** By thresholding the data at high statistical significance, the volume will have high sparsity; the number of elements which are insignificant (NaN, or below threshold) will vastly outweigh those elements which contain numeric, non-zero values.
2. **Smoothness.** As each element shares a contiguous relationship with its neighbours in both the spatial and time dimensions, the volume as a whole, as well as the clusters of significant activity within the volume will have an inherent smoothness.
3. **Unipolarity.** The statistical *t*- and *F*-tests performed on the data only result in positive real numbers.

### 2.3 Porthole

*Porthole* (`porthole.m`) is our toolset for visualisation of spatiotemporal statistics as animated scalp maps. As shown in Figure 2, the main *display window* comprises a two-dimensional cartesian grid system, overlaid with a scalp outline with nose and ears used to indicate head orientation. The user also has the option for a secondary channel coordinate overlay for interpretation in sensor space. Iterating through each time slice, the animation is performed by assigning variations colour to each element in the grid.

#### 2.3.1 Visual components

The display window is also framed by a number of additional objects which provide the necessary contextual information to gain a full appreciation for the data.

The *legend* (bottom left corner, `ph_get_legend.m`) is a general point of reference, summarising the contents of the dataset as a whole. This indicates the type of data being shown, the criteria used to threshold the data, the global maxima and minima within the dataset, and the colour mapping between them. Our default colour mapping transitions from red to yellow, which denotes an increase in statistical significance with a lower bound (Christen et al., 2013).

The *timeline* (right, `ph_get_timeline.m`) gives the user a sense of the overall response and distribution of data along the *t*-axis. The length of the timeline is coloured by extracting the local maxima within each time bin (*non-planar drilling* in Bach et al., 2016). This also serves a similar purpose as the single channel asterisk annotation as shown in Figure 1A, indicating the time windows of statistical significance and associated level of confidence. An arrowhead (`ph_get_arrow.m`)is translated along the timeline with the animation, indicating the current position within the volume and forms a key navigational tool when using the interactive mode.

The *information readout* (top left corner) displays the numerical data relating to the current time slice. Parameters listed are the image index, peri-stimulus time, the local maxima and the size and spatial location of peak local significance, shown in cartesian coordinates. For example, in Figure 2, we show image number 100 of the total 101 images referring to time *t* = 395 ms, with peak significance of *t*-value = 6.0576 spatially located at voxel (18,21) on the *x*-*y* plane.

#### 2.3.2 Importing data

For the visualisations to run independently of the SPM environment, we import these datasets from the universal NIfTI-1 format (.nii) into generic three-dimensional arrays and save in MATLAB format (.mat) via the `nii2ph` function.

**Figure 2.**
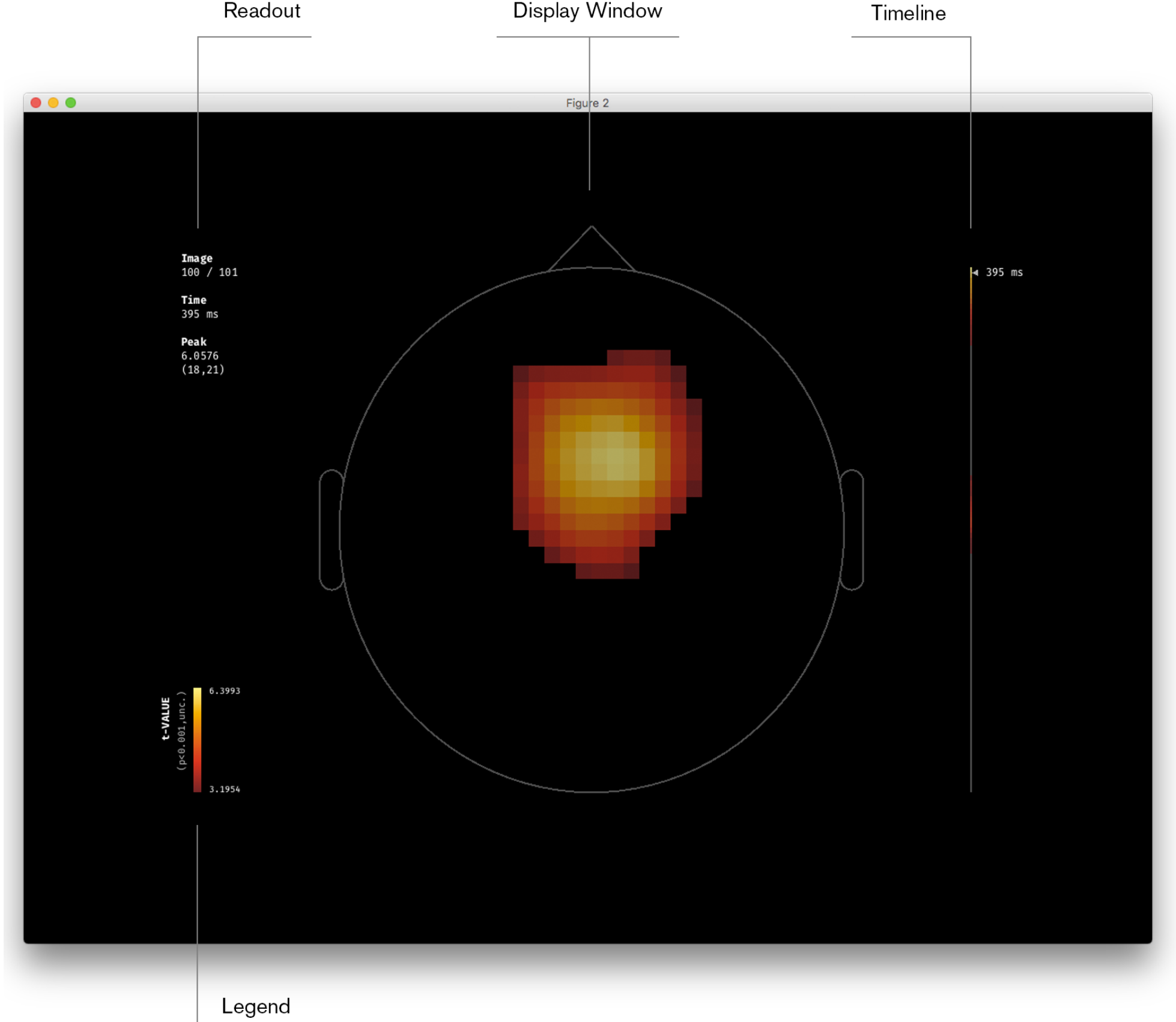
Example frame from *Porthole* visualisation. Scalp map animation is performed by iteratively assigning colours to each element in the display window. The timeline summarises the overall response and indicates current temporal position within the volume. The display window is framed by contextual metadata in the legend and information readout. Animation can also be controlled manually via user interaction. Data shown is a statistical parametric map, illustrating the main effect of surprise, contrasting standard and deviant evoked responses to an auditory oddball paradigm.

#### 2.3.3 Initialising the visual environment

Each dataset also requires specification of metadata and user preferences via graphical user interface (`ph_gui.m`, shown in Appendix A), which are appended to the data structure.

The data type, the *p*-value and correction associated with the SPM thresholding are shown in the legend. By default, the colour mapping is autofit to the global minimum and maximum values within the dataset. The user can also specify their preferred thresholds, which may be useful in defining a common legend to be shared by a series of visualisations, enabling direct comparison between them, or lowering the precision of these extrema.

In order to annotate the timeline and readout, the epoch timing is defined by the sampling frequency and pre-stimulus interval. As the effects of interest may occur late within the epoch, the bounds of the animation loop can be controlled by specifying the desired start and finish time.

The display window can be customised by setting the scalp shape to be circular or oval (0.8 width × 1 height voxel dimensions), with options to overlay channel locations and/or channel labels (`ph_channel_plot.m`) using the 10-10 or 10-20 international standard template for EEG electrode placement (Oostenveld and Praamstra, 2001).

#### 2.3.4 Animating the display window

To perform the animation process, the face colours of each element in the display grid are reset according to the changes in data values read across time slices. The range of colours able to be displayed on-screen are defined in a three-dimensional RGB (or red, green and blue additive) colourspace, packaged as a three element vector or triplet. As such, we are able to perform standard linear algebra operations on this colour space. Should the user wish to display the image sequences at slower speeds, additional frames can be generated by performing a sinusoidal interpolation between the colour triplets assigned to elements (*x*,*y*,*t*) and (*x*,*y*,*t*+1) through vector calculus.

Two different modes are provided for the user to display their datasets in the desired manner; *animation* mode and *interactive* mode. In animation mode, the image sequences are played automatically. Interactive mode pauses the animation and enables the user to iterate between images manually, without interpolation, using the arrow keys. Animation mode is the default setting upon startup and the display mode can be toggled at any time by pressing the space bar.

### 2.4 Stormcloud

*Stormcloud* (`stormcloud.m`) is an extension of the *Porthole* framework used for rendering the full spatiotemporal data as a volume, containing discrete cloud-like clusters with scalp map annotations referring to peaks within those clusters and are intended for publishing in print. *Stormcloud* uses a shared data structure with *Porthole* (Section 2.3.2) and a similar graphical user interface (`sc_gui.m`, shown in Appendix B).

#### 2.4.1 Rendering the volume

To model the data volumetrically (as illustrated in Figure 3), we first create a set of small cubic objects with unitary vertices for each voxel above threshold within the dataset. Rather than mapping colours to the data points using RGB components as in *Porthole*, the voxel face colour is here set to a constant mid-grey and the appearance is instead controlled by manipulating transparency (or opacity) using a fourth component, *alpha*. These alpha values control whether a surface appears fully transparent (α = 0), opaque (α = 1), or at infinitely many levels of semi-transparency (0 < α < 1). By normalising the dataset, we can therefore perform a direct mapping such that the significance level for each voxel is proportional to its transparency. When the collective set of voxels are rendered in this manner, the clusters have a cloud-like appearance, where highly significant elements appear darker, less significant elements are lighter and insignificant voxels are invisible.

The volume is presented in isometric perspective, defined in spherical coordinates as azimuth angle of 45° (rotated in the *x*-*y* plane around the *z*-axis), and elevation angle of 35.264° (i.e. arctan(1/√2), rotated from the *x*-*y* plane toward the *z*-axis). In isometric perspective, the angles between the *x*, *y* and *z*-axes appear equal (120°), providing a level of spatial accuracy which is more directly interpretable than other methods of 3D projection — parallel lines appear parallel, lines of the same length appear to have the same length, and surfaces with the same area appear to have the same area.

However, although we are rendering the volume in three-dimensions, through the printing process we are ultimately presenting the volume on a two-dimension surface from a single viewpoint. As many datasets contain irregular shapes, a single viewpoint can obfuscate some of the data in the background. There are four possible isometric viewing angle to choose from, which we describe in terms of the scalp orientation and which quadrant is closest to the observer; right-posterior, right-anterior, left-anterior, and left-posterior. We recommend rendering the volume from all four of these different viewpoints and selecting those which best articulate the effects to be conveyed.

#### 2.4.2 Annotating the volume

Whilst loading the dataset, *Stormcloud* automatically identifies the discrete clusters of activity within the volume, as well as the number of voxels and points of peak significance within these clusters. Using these criteria as a guide, the author can then specify which clusters they wish to annotate, such that they can be explicitly referred to within the text. Using the `ph_draw_scalp` function, scalp outlines are drawn around the volume at the time points when these peaks occur and a companion two-dimensional scalp map can be displayed next to these points of interest.

The set of scalp maps is obtained from given time indices using the `ph_export_maps` function, which generates high-resolution images akin to the *Porthole* display window and saves them in the working directory. Like the *Porthole* customisation options, scalp shape can be specified as a circle or oval, and custom colour map thresholds can also be adjusted.

## 3. Results

In this section, we present three examples which highlight the strengths of these visualisation tools and possible applications. The experimental effect being analysed is known as Mismatch Negativity (MMN; Näätänen, 1990), which can be simply described as sensory prediction error, the difference between responses to predictable and unpredictable stimuli, or the brain’s response to surprise. The dataset used was provided by Harris et al. (2018) and is freely available at https://figshare.com/s/1ef6dd4bbdd4059e3891. *Porthole* video screen captures corresponding to each of these examples are available in the supplementary materials. Pre-alpha versions of the *Stormcloud* toolset have also been used previously by Garrido et al. (2016), Timmermann et al. (2017), Larsen et al. (2018) and Garrido et al. (2018).

### 3.1 Dataset

#### 3.1.1 Experimental design

EEG was recorded whilst participants listened to auditory stimuli with two overlaid components; Gaussian white noise, which was played binaurally, and an auditory oddball paradigm that used pure tones, played monaurally. The oddball paradigm comprised a sequence of pure sinusoidal tones, 50 ms in duration, presented at 500 ms intervals and occasionally deviating in frequency in 15% of trials (500 or 550 Hz, counter-balanced between blocks). Silent intervals or ‘gaps’ were also embedded within the white noise, which could be singular (90 ms) or repeated (two 90 ms gaps, separated by 30 ms of noise). In a given block, participants were asked to pay attention to these gap stimuli in one ear, ignoring those in the other ear. On a numbered keyboard, they then pressed the number ‘1’ in response to the single gap stimulus and ‘2’ for the double gap. For more information regarding the task, refer to Garrido et al. (2018).

**Figure 3.**
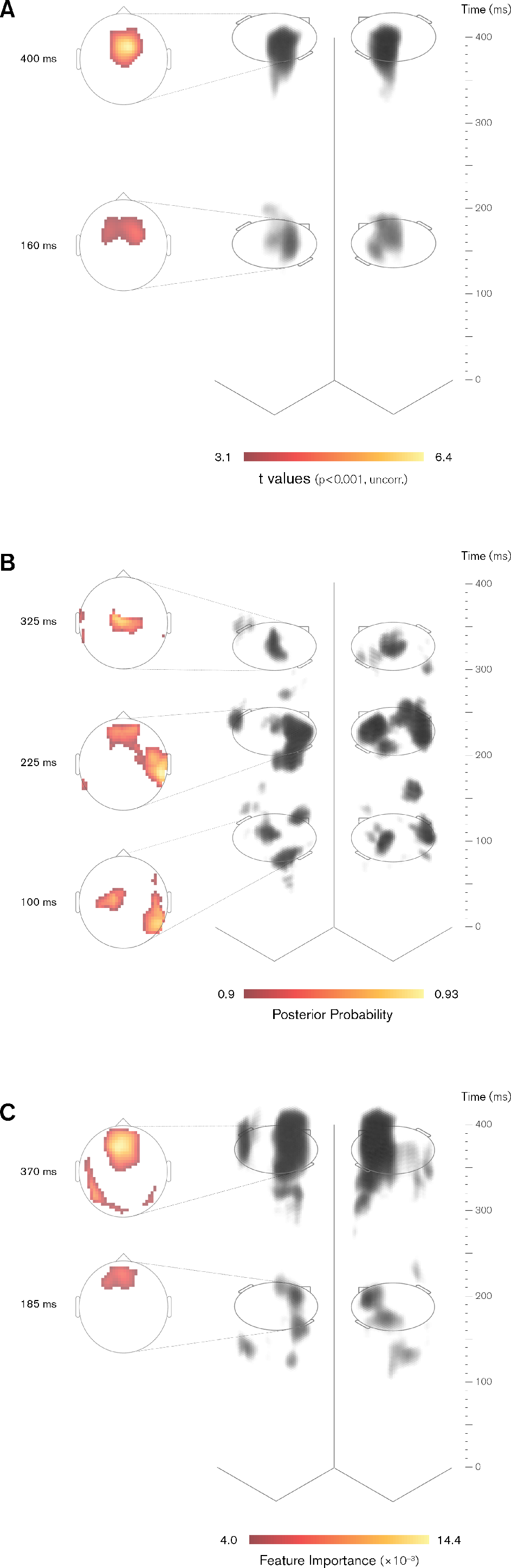
Examples of *Stormcloud* visualisations. 3D spatiotemporal clusters rendered from dual isometric perspectives with spatial dimensions on *x-y* plane, time domain along *z*-axis, and voxel transparency mapped to level of statistical significance. Peaks within clusters of interest are annotated by cross-sectional 2D scalp maps. **(a)** Statistical parametric map — main effect of surprise, computed via *t*-contrast between standard and deviant responses, thresholded at *p* < 0.001 (uncorrected). **(b)** Posterior probability map — evidence that attention boosts the evoked responses to both standard and deviant stimuli, computed via Bayesian model selection and thresholded at 90% posterior probability. **(c)** Machine learning feature importance map — weights obtained from binary support vector machine classification between unattended standard and attended deviant responses, multiplied by grand mean image and thresholded at top 5% highest contribution to model predictions.

#### 3.1.2 Participants

A group of 21 healthy adults with no reported history of head injury resulting in unconsciousness, and no mental or neurological disorders (age 19 to 64 years, mean = 25.00 years, *SD* = 9.83, 12 males) participated in the study. All participants provided written and verbal informed consent and were compensated for their time in accordance with guidelines set by the University of Queensland Human Research Ethics committee.

#### 3.1.3 Data collection and pre-processing

Continuous EEG data were recorded using a 64 channel BioSemi ActiveTwo system (Amsterdam, Netherlands) with electrode placement in accordance with the international 10-10 standard at a sampling frequency of 1024 Hz. Offline signal processing was performed using SPM12 (http://www.fil.ion.ucl.ac.uk/spm/). Data were re-referenced to the scalp average, high-pass filtered using a Butterworth filter with cutoff frequency at 0.5 Hz, downsampled to 200 Hz and low-pass filtered at 40 Hz. Experimental trials were epoched with −100 to 400 ms peri-stimulus window and baseline corrected to the −100 to 0 ms pre-stimulus interval. Trials with signal amplitudes exceeding a ±80 μV threshold were excluded from the analysis and ERPs for each condition were obtained by averaging across trials. Across participants, the mean stimulus count following artefact rejection was 1026.8 standards and 186.0 deviants for the attended condition (15.3% deviants), 1127.3 standards and 200.8 deviants for the unattended condition (15.1% deviants). ERPs were converted to a set of four spatiotemporal NIfTI images per subject, interpolating scalp data into a spatial 32 × 32 matrix for each time bin (101 samples from −100 to 400 ms, 5 ms increments), then smoothed using a Gaussian filter with FWHM 8 × 8 × 8 voxels.

### 3.2 Statistical parametric maps

Using the mass-univariate general linear modelling approach outlined in Section 2.2.1, we analysed these spatiotemporal images using a 2 × 2 design with surprise and attention as factors, specifying four regressors: standard and deviant under attended and unattended conditions. We then computed *t*-contrasts across the group with a cluster forming threshold at *p* < 0.001 (uncorrected). As illustrated in Figure 3A, we found two large clusters for the main effect of surprise, peaking at 160 ms (cluster-level *p* = 8.76 × 10^−4^, FWE corrected) and 400 ms (cluster-level *p* = 6.99 × 10^−4^, peak-level *p* = 6.51 × 10^−5^, FWE corrected), both located in fronto-central scalp regions. There were no significant main effects of attention or interactions between surprise and attention.

### 3.3 Posterior probability maps

Another means of analysing spatiotemporal images is through probabilistic Bayesian inference. By placing a prior assumption on the general effects we expect to see in the data, we can then update the likelihood of this assumption being true, given the data we actually observe. Such testable assumptions can range from a simple null effect (or null model that is reminiscent of the frequentist approach in used SPMs), to more sophisticated alternative assumptions (or models) about what has generated the data. This is formally known as Bayesian model selection (BMS) and is particularly useful for comparing computational models at the group level. In this example, we focus on one model in particular, which proposes that attention boosts the evoked responses to both standard and deviant stimuli. This implies that, across the four conditions, activity evoked by unattended standard stimuli has the lowest amplitude, attended deviant is greatest, with attended standard and unattended deviant approximately equivalent (Garrido et al., 2018). The probability map supporting this model, shown in Figure 3B (also described in Harris et al., 2018) was thresholded at 90% posterior probability.

At each voxel, we can compute the model evidence, or the likelihood of the observed data from individual participants under the assumption that the model is true. The evidence for the model can then be converted to a posterior probability map at the group level. For more detail on this methodology behind these computations, refer to Harris et al. (2018) and Rosa et al. (2010).

### 3.4 Machine learning weight maps

Machine learning methods are being increasingly applied to neuroimaging data as a means of decoding multivariate patterns of neural activity associated with a given experimental condition. In this example, spatiotemporal images obtained from each subject were used as inputs for a machine learning model using the Pattern Recognition for Neuroimaging Toolbox (PRoNTo; Schrouff et al., 2013), such that each individual voxel was considered a learning feature. In contrast with the mass-univariate approach, these multivariate techniques instead jointly consider pairwise similarities between all voxels in the image (kernel method; Schölkopf and Smola, 2000).

Using the Support Vector Machine algorithm (SVM; Burges, 1998), we trained a model to classify images according to the labels ‘unattended standard’ and ‘attended deviant’ as a proof of concept, noting that the differences in ERP were maximised between these two conditions. Considering each image as a datapoint in a high-dimensional space, SVM generates a hyperplane which best separates the two conditions by maximising the margin between them (we use the default soft-margin hyper-parameter, *C* = 1). This hyperplane then serves as a binary decision boundary. As part of this training process, each voxel within the image is assigned a weighting, which represents the relative contribution of that voxel toward classification. Predictions for new test images are then made via multiplication with the weight map, returning a signed scalar value indicating which side of the hyperplane the image lies (in this instance, positive and negative values are classified as ‘standard’ and ‘deviant’, respectively).

To assess the overall performance of the model, we employed a 10-fold cross-validation scheme (Hastie et al., 2001), which partitions the images into 10 subsets, then iteratively assigns 90% of the images for training and the remaining 10% for testing the model. Overall, this model was able to discriminate between the two experimental conditions with 80.95% total accuracy (*p* = 0.001, permutation test), 85.71% (*p* = 0.002) class accuracy for ‘standard’ and 76.19% (*p* = 0.025) for ‘deviant’ responses.

Although it is commonplace to inspect the resulting weight maps and intuit which points in space and time are driving the model performance, direct interpretation is not straightforward (Haufe et al., 2014). As all voxels contribute to the predictions, we cannot fully comprehend the interactions between them on face value. For example, given that the original images and the weight map both contain signed values, a negative weight applied to a negative feature has a net positive influence on the model prediction. Similarly, a large weight can also be applied to a small feature, or vice versa, although when the data has adequate signal-to-noise ratio, a large weight assignment is unlikely to be a false positive (Schrouff and Mourão-Miranda, 2018).

To better understand which features informed ‘standard’ predictions, we computed the element-wise multiplication between the mean weight map (averaged across folds) and the grand mean image (averaged across participants). We refer to the resulting as an ‘importance map’. When thresholded at the top 5% largest values, as shown in Figure 3C, this importance map summarises 88.79% of the total contributions toward a ‘standard’ classification, with the remaining voxels likely suppressing noisy or irrelevant information. It is worth noting that the mass univariate and multivariate approaches, displayed in Figure 3A and 3C, respectively, are remarkably consistent both in terms of the temporal and spatial extent of the significant clusters. However, as expected, the multivariate approach is much more sensitive than the univariate approach, thereby revealing a greater number and extent of clusters.

## 4. Discussion

M/EEG data can be represented volumetrically as a three-dimensional space-time construction. Using computational modelling techniques such as statistical parametric mapping, we are able to analyse the rich spatiotemporal data as a whole, rather than selecting subsets of channels and/or time windows by a priori assumption regarding the effects we expect to see. The resulting images output from this modelling contain a set of discrete internal clusters with an inherent smoothness. Although the conventional methods of M/EEG visualisation each have their own strengths, they are only able to present a subset of the data, focusing primarily on articulating one of the space-time components. These visualisation techniques can be categorised as single channel time series, topographic scalp maps and orthographic cross-sections. Single channel plots, obtained from a given point in space on the *x*-*y* plane, have a high temporal resolution, but little to no spatial information. Conversely, topographic scalp maps extracted from a given time point *t*, have very high spatial resolution, but do not capture temporal information. Although a sequence of scalp maps can be presented in grid format, these are often grossly downsampled, hence dramatically comprising the temporal resolution. Orthogonal cross-sections are able to provide high spatiotemporal resolution with respect to a given point (*x*,*y*,*t*), but are unable to provide a full appreciation of the cluster shape and extent without interactive user navigation.

In this paper, we introduced *Porthole* and *Stormcloud*, a set of visualisation tools which enable in print and on screen visualisation of M/EEG statistics across the whole spatiotemporal topography in a more comprehensive, informative and efficient manner using a combination of two-dimensional scalp animation and three-dimensional rendering techniques. In addition, we suggest that these methods offer improved resolution of all three space-time components and also demonstrate their intended application in the context of our own empirical work.

*Porthole* centres around two-dimensional topographic scalp animation, which can be described as an iterative slicing operation across the spatiotemporal volume at each individual time point. However, passively watching an animation necessitates that the viewer retains information in working memory, which may not suffice in gaining a full appreciation of the data — they must be aware of the current time point within the epoch, the spatial effects that have occurred previously, and potentially anticipate the spatial effects that will occur in the future. As a countermeasure, we frame the display window with a dynamic timeline, colour coded by the local maxima extracted from each scalp map. This is akin to the asterisk annotation from the single channel plots shown in Figure 1A, indicating the time windows of statistical significance and associated level of confidence. We also provide the option for user interaction to pause the animation sequence and navigate the volume manually. Collectively, these methods provide high resolution in both spatial and temporal components of the data.

Kristensson et al. (2009) suggest that space-time cube visualisation systems similar to those presented here are more effective at communicating complex spatiotemporal information. In that study, users answered a series of questions by interpreting spatiotemporal data from either a static two-dimensional map with time-related annotations, or a three-dimensional space-time cube representation which could be navigated interactively. Whilst the two-dimensional map provided significantly lower error rate for questions involving interpretation of the whole space at a single time point, the space-time cube resulted in halved response time (down from 121 to 60 seconds) for more complex questions involving the integration of information across all space and time.

*Stormcloud* presents the spatiotemporal data volumetrically, rendering clusters of statistical significance with a cloud-like effect by scaling the opacity of individual voxels according to the level of significance. As these clusters may have irregular shapes and distributions, rendering the topography in three dimensions may present problems when depicting the volume from a singular viewpoint, as some portion of the data can be obfuscated in the background. As such, we depict the volume in four orthogonal views using common axes and select the pair which best articulates the overall topography. We also annotate the main clusters of interest using companion topographic scalp maps at time points of peak statistical significance. As noted by Fuchs and Hauser (2009), a hybrid of multiple visualisation methods applied to different subregions of the volume can be especially useful in providing the necessary context for correct understanding. *Stormcloud* has moderate spatial and temporal resolution for the volume as a whole, and high spatial resolution at the specified time points.

Whilst these tools were optimised for use with statistical parametric maps (SPM) and our design process was informed by the underlying properties of M/EEG neuroimaging data, we also demonstrated alternate applications with Bayesian posterior probability maps and feature importance maps derived from machine learning weights. In principle, we envision that many, if not all of these methods are transferrable to essentially any spatiotemporal image.

## Appendices

**Appendix A.**
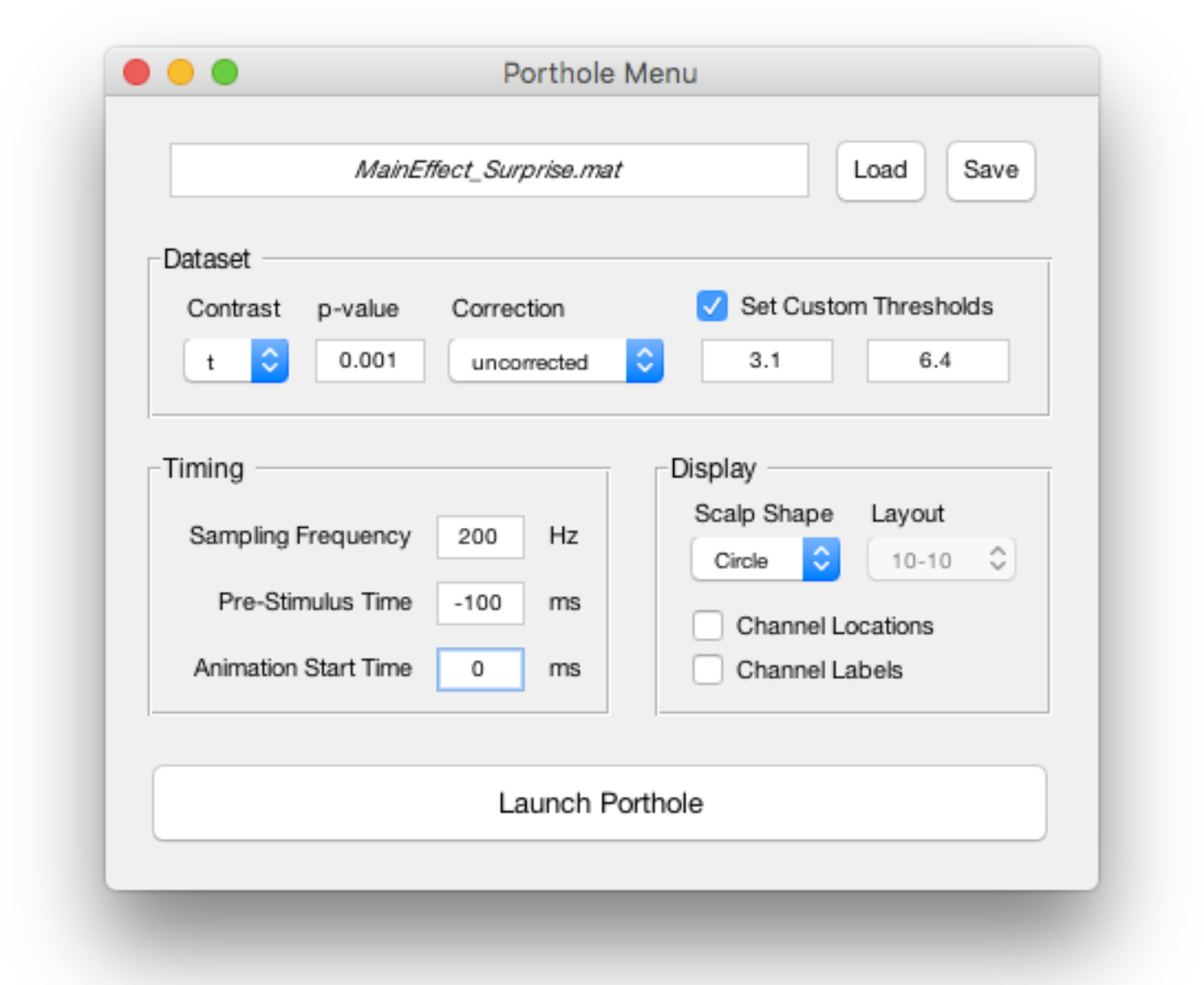
Graphical user interface for specifying *Porthole* animation parameters and metadata for annotating the display.

**Appendix B.**
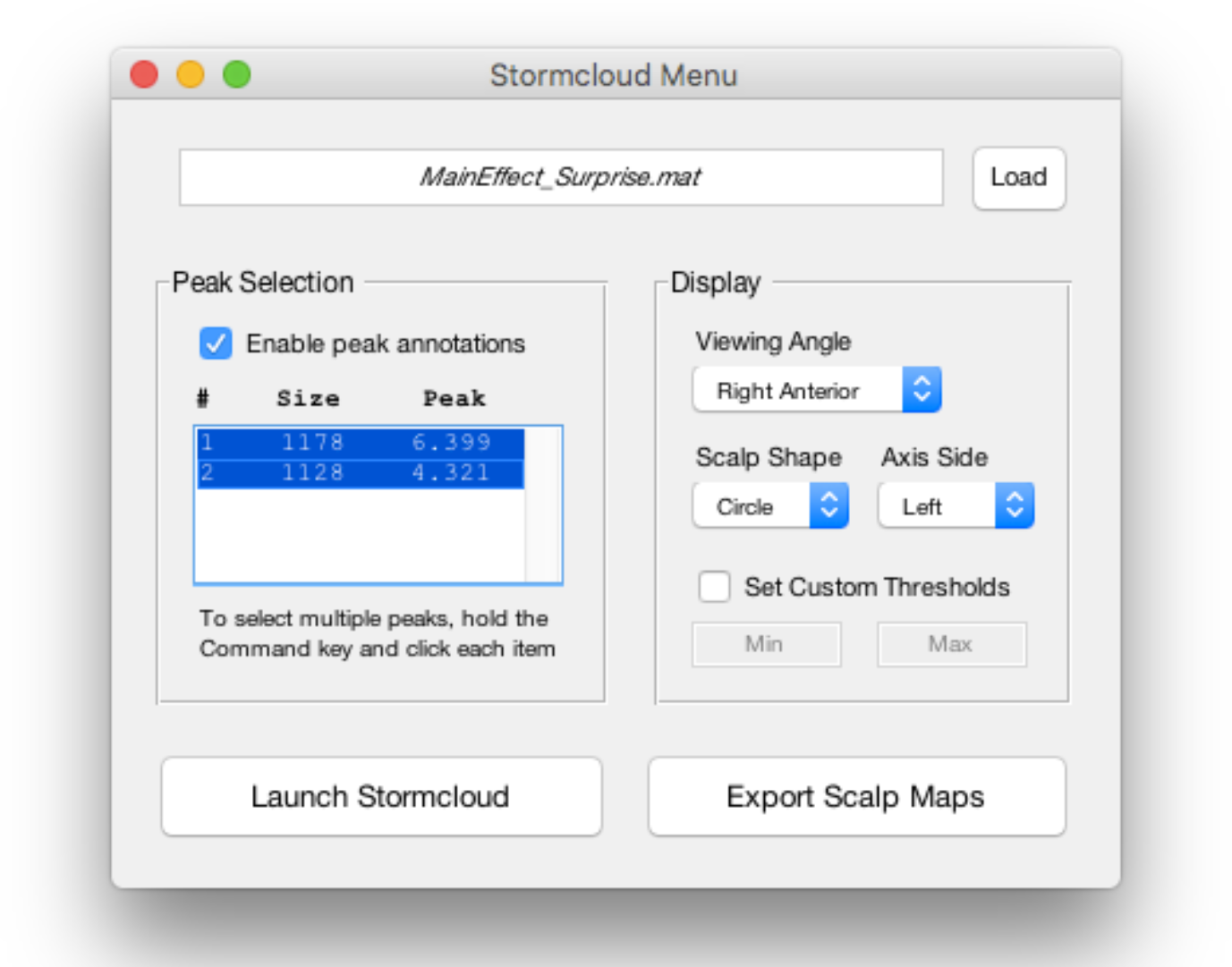
Graphical user interface for specifying *Stormcloud* preferences for volume orientation and cluster annotation.

## Acknowledgements

This work was supported by the Australian Research Council Centre of Excellence for Integrative Brain Function (ARC Centre Grant CE140100007), a University of Queensland Fellowship (2016000071) and a Foundation Research Excellence Award (2016001844) to MIG. We would like to thank Tyler Hobson for discussions on computer graphics methods, Clare Harris for providing data, as well as Veronika Halász, Kit Melissa Larsen, Ilvana Dzafic, Jessica McFadyen and Chase Sherwell for providing feedback on the functionality of earlier versions of the toolbox.

